# Cross-phosphorylation of RR06 by the non-cognate kinase VncS activates the adhesin CbpA in *Streptococcus pneumoniae*

**DOI:** 10.64898/2026.07.15.738638

**Authors:** Si-Yin Tan, Justin J. Zik, Ye-Yu Chun, Kai Sen Tan, Lok-To Sham

**Affiliations:** Infectious Diseases Translational Research Programme and Department of Microbiology and Immunology, Yong Loo Lin School of Medicine, National University of Singapore, Singapore

## Abstract

*Streptococcus pneumoniae* establishes asymptomatic nasopharyngeal colonization before progressing to invasive disease, a process that requires precise spatiotemporal regulation of surface adhesins. Using a barcoded capsule-switch mutant library and randomly barcoded transposon sequencing (RB-TnSeq), we identified determinants of pneumococcal adhesion to the human alveolar epithelial cell line A549. The capsule masked surface adhesins and reduced binding across all 84 serotypes tested. The transposon screen revealed that overexpression of *vncS*, the sensor kinase of the two-component system TCS10, markedly increased adhesion, whereas disruption of its cognate response regulator VncR had no detectable effect. Transcriptomic profiling and genetic epistasis established that the adhesin CbpA, known to be regulated by TCS06, is the effector of VncS-driven adhesion. VncS activated *cbpA* expression through the non-cognate response regulator RR06, independently of VncR. Phos-tag gel analysis demonstrated that VncS cross-phosphorylates RR06 *in vivo*, and that the cognate kinase HK06 limits this crosstalk by acting primarily as a phosphatase toward RR06. Substitution of the conserved S241 residue of HK06 increased RR06∼P levels and upregulated *cbpA*. By contrast, deleting *vncS* and *hk06* reduced RR06 phosphorylation to nearly undetectable levels. These findings resolve a long-standing mechanistic controversy regarding RR06 activation and uncover a potentially unrecognized role for VncS in regulating pneumococcal host cell adhesion.

**SIGNIFICANCE:** *Streptococcus pneumoniae i*s a leading cause of pneumonia, and its ability to colonize the human airway depends on surface adhesins. Several two-component signaling systems are known to regulate adhesin expression, but how their outputs are integrated remains incompletely understood. Here, we identify determinants of pneumococcal adhesion to human alveolar epithelial cells at both the serotype and genotype levels. We show that the histidine kinase VncS cross-phosphorylates the non-cognate response regulator RR06 to activate the adhesin CbpA, whereas the cognate kinase HK06 primarily acts as a phosphatase to restrain this crosstalk. These findings resolve a long-standing controversy regarding RR06 activation and reveal a regulatory node that links two signaling systems to a single virulence output.

## INTRODUCTION

*Streptococcus pneumoniae* (the pneumococcus) is a leading cause of community-acquired pneumonia. According to the Global Burden of Disease Study 2023, lower respiratory infections (LRIs) were responsible for an estimated 2.50 million deaths and 98.7 million disability-adjusted life years (DALYs) worldwide [1]. Pneumococcus alone caused over 600,000 deaths and 58.3 million lower respiratory infections (∼24%), the largest share among all etiological agents in that study [1]. The burden is borne disproportionately by children under five and adults over 70, and significant disparities persist in low-income settings [1]. Despite the availability of polysaccharide-based conjugate vaccines [2], pneumococcal disease continues to exact a massive global toll, underscoring the need to better understand the molecular mechanisms that govern pneumococcal virulence and host interactions.

The pathogenesis of invasive pneumococcal disease (IPD) often begins with nasopharyngeal colonization [3]. *S. pneumoniae* can persist in the nasopharynx asymptomatically for weeks to months [4,5]. Yet this colonization serves not only as a reservoir for lower respiratory tract infections when the host immune system is compromised, such as after influenza infection, but also as a source of horizontal transmission among individuals [3,4,6]. Central to pneumococcal colonization is the capsular polysaccharide (CPS) [7]. The capsule shields the pneumococcus from complement deposition and opsonophagocytic clearance by host neutrophils [8]. It is typically negatively charged, which is thought to repel mucus, thereby facilitating access to the underlying epithelial layer [9]. However, the capsule simultaneously occludes surface-exposed adhesins from host receptors. To expose these adhesins, *S. pneumoniae* employs several strategies. First, phase variation leads to heterogeneity in capsule synthesis within the population, thereby enhancing nasopharyngeal colonization [10]. Capsule shedding mediated by LytA, particularly prominent in serotype 3, allows the thick capsule to be locally released, unmasking adhesins at the epithelial surface [11]. Additionally, RNA thermosensors detect temperature shifts from the nasopharyngeal cavity to the bloodstream, suppressing capsule expression at lower temperatures to promote colonization [12]. The capsule promoter is also transcriptionally controlled, mostly through a conserved 37-base-pair cis-regulatory element called the 37-CE [13–15]. These regulatory mechanisms ensure that surface adhesins are unmasked in the nasopharyngeal niche while maintaining the ability to restore full capsule production during systemic infections.

Among the adhesins that mediate host-pathogen interaction, choline-binding protein A (CbpA, also known as PspC) plays a major role in mediating adherence and invasion of the nasopharyngeal epithelium. It binds directly to the polymeric immunoglobulin receptor (pIgR) [16,17], facilitating translocation across nasopharyngeal epithelial cells for invasive infections [17]. In addition, CbpA recruits human complement factor H and downregulates the alternative complement pathway [18,19]. The expression of CbpA is controlled by the two-component regulatory system TCS06 [20–22]. Given its role in pathogenesis, CbpA is a potential vaccine target [23,24]. *S. pneumoniae* also produces additional adhesins that contribute to colonization. Neuraminidase A (NanA) cleaves terminal sialic acid residues from host glycoproteins and is implicated in promoting biofilm formation [25]. The underlying glycans are further processed by BgaA, StrH, and EndoD, thereby exposing host-cell receptors [26,27]. The sugars cleaved by these enzymes are also utilized after import via various transporters [27]. Other adhesins include pneumococcal adherence and virulence protein A (PavA), which bridges pneumococcal surface proteins with extracellular fibronectin. The coordinated surface expression of these adhesins in appropriate host niches is essential for colonization and disease progression [6,9].

Understanding the mechanism governing pneumococcal adhesion is therefore crucial for elucidating pathogenesis. Previously, we infected the human nasal epithelial cells (hNECs) and bronchial epithelial cells (hBECs) with a collection of barcoded isogenic capsule-switched mutants [28]. This library comprises 84 out of the 108 known serotypes [29]. Here, we continue our investigation using a random-barcoded transposon (RB Tn) library to systematically identify determinants of bacterial adhesion to the human alveolar epithelial cell line A549 [30]. This unbiased approach unexpectedly revealed the crosstalk between the histidine kinase VncS (in TCS10) and the response regulator of TCS06 (RR06). By RNA sequencing and Phos-tag phosphorylation assays, we demonstrate that overexpression of *vncS* increased bacterial adhesion to host cells through upregulation of *cbpA*, independently of its cognate response regulator VncR. This upregulation is due to phosphorylation of RR06 by VncS, which is counteracted by the phosphatase activity of the cognate histidine kinase HK06. These findings reconcile the longstanding mechanistic controversy of *cbpA* regulation and reveal a previously unrecognized role for VncRS in pneumococcal virulence.

## RESULTS

### Capsule impedes binding of *S. pneumoniae* to alveolar epithelial cells

To investigate the role of CPSs in attachment to alveolar cells, we introduced the barcoded capsule-switched mutant library into A549 cells (**Fig. S1**). Although A549 cells do not form an air-liquid interface (ALI), they are a well-established respiratory epithelial model that avoids the donor-to-donor variability common in primary cell cultures [28]. Pneumococci were inoculated into the A549 cell culture, centrifuged, and allowed to co-culture for three hours. Unbound pneumococci were recovered by aspirating the culture medium. The attached *S. pneumoniae* was collected by trypsinizing adherent A549 cells. The bound and unbound fractions were plated, and the DNA barcodes were sequenced. After measuring the relative abundance of pneumococcal serotypes with barcode sequencing, we confirmed that the unencapsulated (Δ*cps*) strain bound substantially more to A549 cells (**Fig. S1**). This result shows a strong correlation between the two biological replicates (ρ = 0.9281, P < 0.0001), validating the reproducibility of the barcode sequencing (Bar-seq) approach. To validate these findings, we individually tested 12 representative serotypes that span a wide range of A549 cell adherence in the binding assay. The individual validation results correlated well with the Bar-seq screen (**Fig. S1**). Δ*cps*, serotypes 6C, 33C, and 15C strains bound strongly to A549 cells. Serotypes 36, 6A, 9L, and 22F strains exhibited intermediate binding. Finally, serotypes 19A, 15A, 31, and 2 cells bound relatively poorly compared with the others (**Fig. S1**). These results demonstrate that although capsular serotype influences host cell attachment, the capsule itself is a barrier to adhesion, likely because it blocks adhesins from reaching host cell receptors.

### Overexpression of *nanA* and *cbpA* increases adhesion to A549 cells

To investigate the mechanism of pneumococcal adhesion, we inactivated capsule production by deleting the initiating phosphoglycosyltransferase *cpsE*. Next, we transformed the resulting strain with genomic DNA from a random-barcoded transposon (RB Tn) library containing ∼600,000 unique transposon mutants [30]. Constructing the Tn library in an unencapsulated strain also reduces the bottleneck inherent in library screens by allowing a greater proportion of mutants to be recovered in the bound fraction (**Fig. S1**). It may also expose surface adhesins that engage A549 cells (**Fig. 1A**). After collecting the Tn mutants, we infected A549 cells using the same procedure used to enumerate the capsule-switch mutants and sequenced the DNA barcodes. Bar-seq analysis revealed pronounced enrichment of Tn insertions upstream of two known adhesin loci in the bound fraction. The first locus comprises *spd_1505* and *spd_1506*, located immediately upstream of *nanA*. Another region is upstream of *cbpA*, containing *spd_2018*, *hk06*, *rr06*, and *spd_2021* (**Fig. 1**). Since the barcoded Tn was designed to have a constitutive promoter but lacks a transcriptional terminator, we hypothesize that these Tn insertions drive readthrough expression of the downstream adhesin (**Fig. 1**). Indeed, strand-orientation analysis indicates that Tn insertions are enriched in the same orientation as the transcription of the downstream *nanA* and *cbpA*, respectively (**Fig. 1**). These results suggest that the enrichment of insertions in the bound fraction arises from polar overexpression of *nanA* and *cbpA*, rather than from loss of function of the corresponding genes.

**Figure 1.**
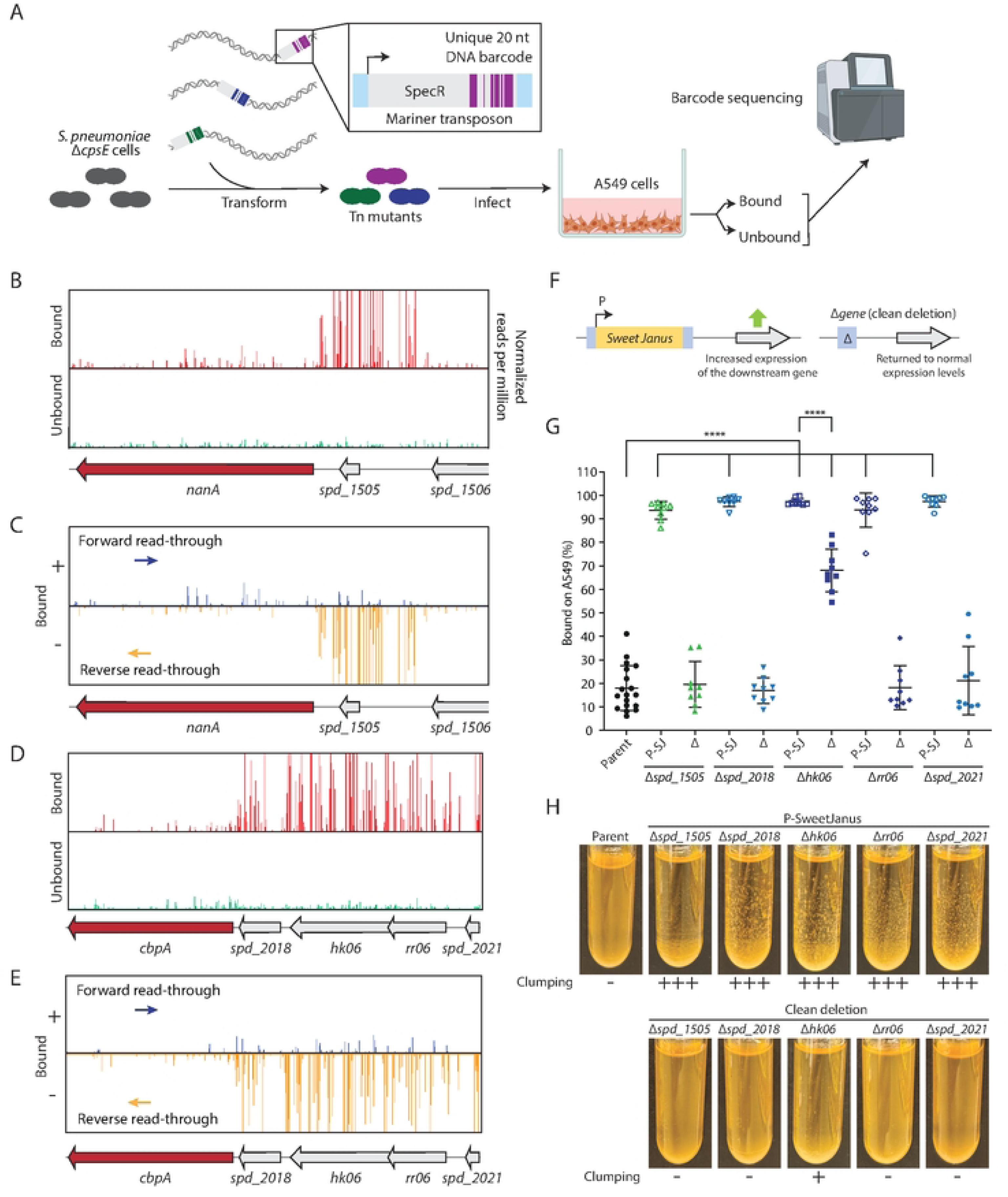
A barcoded transposon screen identifies genes that promote pneumococcal adhesion to A549 alveolar epithelial cells. (**A**) Schematic of the barcoded transposon (Tn) screen. A random-barcoded Mariner transposon library was transformed into unencapsulated *S. pneumoniae* D39 Δ*cpsE*, inoculated onto A549 cell monolayers, and separated into unbound and bound fractions by aspiration and trypsinization, respectively. DNA barcodes were sequenced to determine the relative abundance of transposon insertions in each fraction. (**B**) Genomic tracks showing normalized transposon insertion reads (reads per million) in the bound (top) and unbound (bottom) fractions at the *nanA-spd_1505-spd_1506* locus. (**C**) Strand-resolved insertion plots at the *nanA* locus. Orange bars, insertions on the reverse strand; blue bars, insertions on the forward strand. (**D**) Genomic tracks showing normalized transposon insertion reads in the bound (top) and unbound (bottom) fractions at the *cbpA-spd_2018-hk06-rr06-spd_2021* locus. (**E**) Strand-resolved insertion plots at the *cbpA* loci, separating insertions oriented to drive forward (blue) or reverse (orange) read-through into the adjacent gene. Enrichment of insertions in the bound fraction is consistent with constitutive promoter-driven overexpression of the downstream adhesins. (**F**) Schematic of the validation strategy. Insertion of the Sweet Janus cassette (P-SJ) into a gene increases downstream gene expression via its constitutive promoter. Subsequent deletion of this cassette removes the promoter, returning expression to wild-type levels. (**G**) Quantification of A549 cells adhesion (% bound) for strains carrying the P-SJ cassette or the corresponding clean deletion (Δ) at *spd_1505*, *spd_2018*, *hk06*, *rr06*, and *spd_2021*. Each data point represents an independent biological replicate. Bars show mean ± standard deviation (SD). ****, *P* < 0.0001 by one-way ANOVA with Tukey’s post-hoc test. (**H**) Overnight culture morphology of the SJ-disrupted and clean deletion strains. Self-clumping phenotype was observed after 14 to 18 hours of incubation at 37 °C in 5% CO_2_. +++ denotes strong clumping; + denotes weak clumping; − denotes no clumping. The experiment was conducted three times with similar results.

To test this hypothesis, we reproduced the potential polar effects of the Tn insertions by introducing the Sweet Janus (SJ) cassette into these genes [31]. Like the Mariner Tn, the SJ cassette includes a promoter that drives downstream gene expression and counter-selection markers (*sacB* and *rpsL*^+^) to subsequently generate clean, promoterless deletion mutants (**Fig. 1**). When the SJ cassette was introduced into *spd_1505*, *spd_2018*, *hk06*, *rr06*, and *spd_2021*, the resulting mutants showed markedly increased binding to A549 cells (**Fig. 1**). By contrast, after the SJ cassette was deleted, the corresponding clean deletion strains exhibited binding levels comparable to the progenitor Δ*cps* strain. We also observed that cells carrying the SJ cassette clumped in overnight cultures, a corroborating phenotype reflecting enhanced bacterial surface stickiness and serving as a surrogate assay for A549 cell adhesion (**Fig. 1**). This phenotype reverted upon deletion of the SJ cassette, indicating that the increased binding observed in the SJ-disrupted strains is attributable to polarity affecting downstream adhesins rather than to loss of function of the disrupted gene itself. A notable exception is Δ*hk06*, which retained modest but significantly elevated adhesion (**Fig. 1**). This result was investigated further in the latter part of this study. Next, we tested whether the phenotype is caused by *cbpA* overexpression. First, we introduced the SJ cassette at each locus, but in a Δ*cbpA* background. This change abolished clumping in all strains (**Fig. S2**), demonstrating that CbpA is necessary for the increased adhesion and clumping phenotypes. Quantitative RT-PCR (qRT-PCR) of the *cbpA*^+^ strains showed that SJ insertion in *spd_2018*, *hk06*, or *rr06* elevated *cbpA* transcript levels more than 10-fold relative to their progenitor strain and the corresponding clean deletions (**Fig. S2**). To examine the contribution of NanA to adhesion, we overexpressed a panel of active-site mutants (R328A, E609A, E609Q, Y714F, and E609Q/Y714F) in the Δ*spd_1505*::P-SJ background [32]. All catalytically dead NanA variants retained the ability to induce clumping, whereas deletion of *nanA* abolished it (**Fig. S2**). These results indicate that the presence of NanA, rather than its neuraminidase activity, is required for enhanced bacterial adhesiveness.

### Deletion of HK06 elevates *cbpA* expression and increases adhesion to A549 cells

Unlike other Tn-seq hits mapped upstream of *cbpA*, Δ*hk06* retained a modest increase in binding to A549 cells compared with its parent strain (**Fig. 1**). Since TCS06 is known to regulate *cbpA* expression [20,22], we wondered whether Δ*hk06* altered *cbpA* transcript levels. qRT-PCR showed that *cbpA* expression was elevated approximately 1.5-fold in Δ*hk06* relative to the progenitor strain and was restored to original levels upon ectopic complementation (Δ*hk06* // P-*hk06*) (**Fig. 2**). In contrast, deletion of *rr06* did not result in a detectable reduction in *cbpA* levels (**Fig. 2**), suggesting that residual *cbpA* transcription does not depend on RR06. RNA sequencing of Δ*hk06*, Δ*hk06* // P-*hk06*, and their parent strains revealed very few differentially expressed genes (**Fig. S3**), suggesting that under these experimental conditions, TCS06 may not be activated or that other mechanisms compensate for the loss of *hk06*. Another adhesin, *pspA*, is also known to be regulated by TCS06 [22]. To eliminate the possibility that *pspA* is involved in the difference in attachment in the Δ*hk06* background, we measured the *pspA* transcript level and deleted *pspA*. Indeed, there were no observable changes in *pspA* expression in the Δ*hk06* mutant (**Fig. S3**). Furthermore, inactivating *pspA* did not abolish clumping (**Fig. S3**), suggesting that *pspA* is not a downstream effector of HK06 under our experimental conditions.

**Figure 2.**
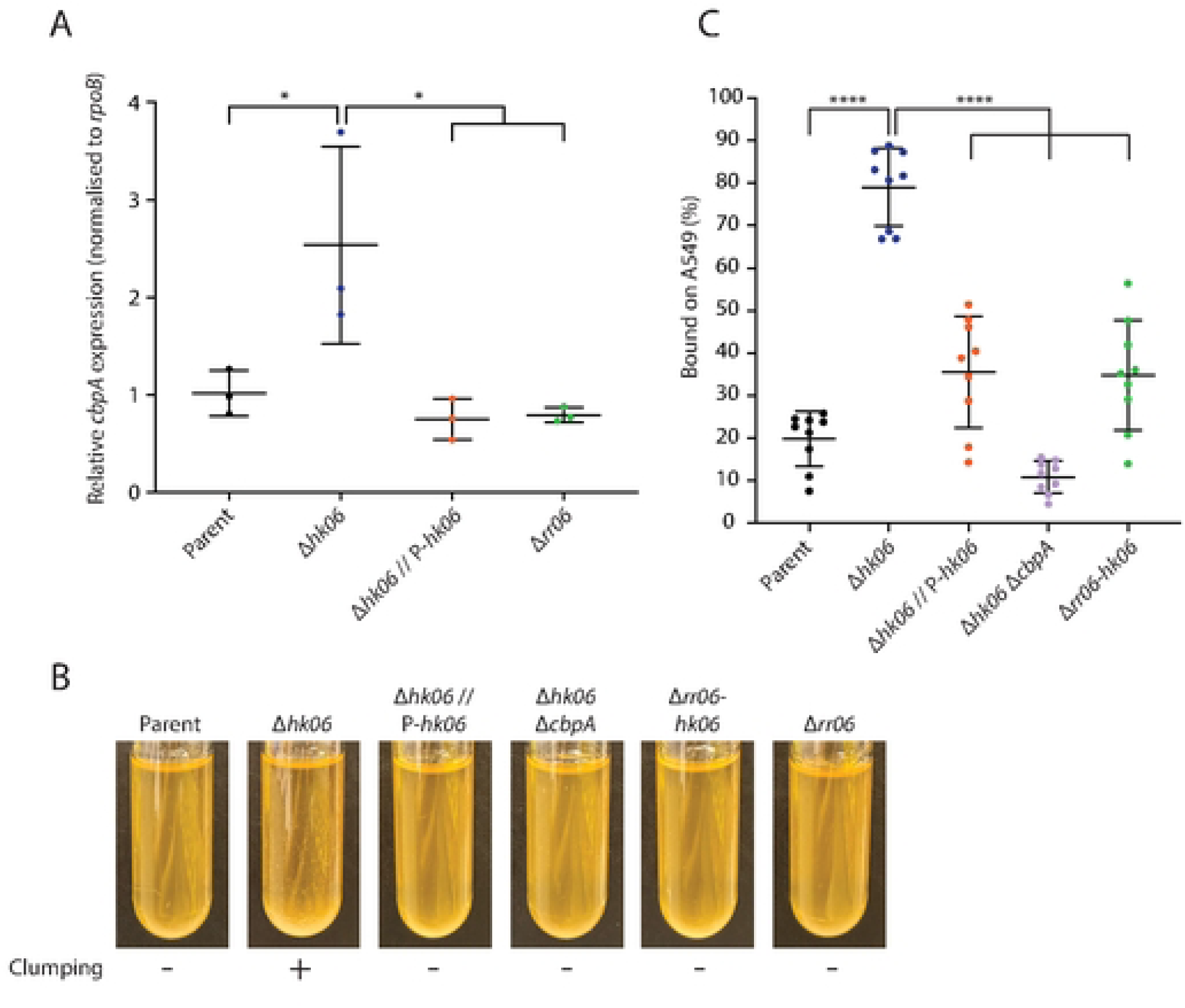
Inactivating *hk06*, but not *rr06*, increases *cbpA* expression. **(A)** Relative *cbpA* transcript levels, measured by qRT-PCR and normalized to *rpoB*, in the parent (HMS0002, D39W *rpsL1* Δ*cpsE*), Δ*hk06* (NUS4966), Δ*hk06* // P-*hk06* (NUS5169), and Δ*rr06* (NUS4967) strains. Each data point represents an independent replicate. Bars show the means from the biological replicates ± SD. *, *P* < 0.05 by one-way ANOVA with Tukey’s post-hoc test. **(B)** Quantification of A549 cells adhesion (% bound) for the indicated strains (HMS0002, NUS4966, NUS5159, NUS6694 [Δ*hk06* Δ*cbpA*], and NUS7202 [Δ*rr06* Δ*hk06*]). Each data point represents an independent replicate. Bars show mean ± SD. ****, *P* < 0.0001 by one-way ANOVA with Tukey’s post-hoc test. **(C)** Overnight culture of the indicated strains. Clumping observed in Δ*hk06* is abolished by complementation with *hk06*, deletion of *cbpA*, or deletion of *rr06*, suggesting that HK06 negatively regulates CbpA-dependent adhesion through RR06. The experiment was conducted three times with similar results.

Corroborating the qRT-PCR data, Δ*hk06* formed visible clumps in overnight broth cultures, while ectopic complementation of *hk06* and co-deletion of *cbpA* each abolished clump formation (**Fig. 2**). Deletion of the entire TCS06 locus (Δ*rr06*-*hk06*) also abolished clumping (**Fig. 2**), indicating that RR06 is still required, perhaps for activating *cbpA* expression. Again, pneumococcal adhesion assays in A549 cells confirmed a strong correlation between the clumping phenotype and host cell binding (**Fig. 2**). Complementation with *hk06* reduced binding by approximately 2-fold relative to Δ*hk06*, to a level similar to that of the progenitor strain, likely reflecting residual polar effects downstream of *cbpA*. Deleting *cbpA* and *hk06* simultaneously reduced binding ∼7-fold relative to Δ*hk06* alone, to levels indistinguishable from the progenitor strain, establishing that CbpA is the primary adhesin responsible for the elevated binding phenotype. Consistent with this, the Δ*rr06*-*hk06* double deletion showed reduced adhesion (**Fig. 2**). Together, these results indicate that HK06 negatively regulates *cbpA* expression through its cognate RR06.

### Overexpression of VncS, but not its cognate RR VncR, promotes adhesion

The Tn-seq screen revealed a second adhesion-associated locus mapped to the TCS10 operon (*vncR-vncS-fba*) (**Fig. 3**). The Tn insertions are enriched upstream of *vncS*, within the coding sequence of its cognate response regulator *vncR*. These insertions were significantly enriched in the bound fraction and were predominantly found in the same direction as the transcription of the operon (**Fig. 3**). This observation is consistent with readthrough transcription, which increases the expression of downstream *vncS*. Notably, no insertions were recovered in the *fba* open reading frame. *fba* encodes fructose-1,6-bisphosphate aldolase, an essential glycolytic enzyme that catalyzes the conversion of fructose-1,6-bisphosphate to dihydroxyacetone phosphate (DHAP) and glyceraldehyde-3-phosphate (G3P) [33]. To distinguish whether the enrichment was caused by Δ*vncR* or by overexpression of *vncS*, we introduced P-SJ cassettes into *vncR,* followed by clean deletion. Introducing the P-SJ into *vncR* caused a ∼3-fold increase in A549 binding and clumping in overnight cultures (**Fig. 3**). The phenotype was corrected by the clean Δ*vncR* deletion, confirming that the observed enrichment was due to transcription readthrough into downstream genes. In contrast, inserting the P-SJ cassette at *vncS* did not increase adhesion to A549 cells. Instead, it slightly reduced binding below that of the clean Δ*vncS* deletion strain (**Fig. 3**). Furthermore, the Δ*vncS*::P-SJ strain did not clump (**Fig. 3**). This phenotypic divergence between Δ*vncS*::P-SJ and Δ*vncR*::P-SJ suggests that *vncS* is necessary for enhanced A549 cell binding.

**Figure 3.**
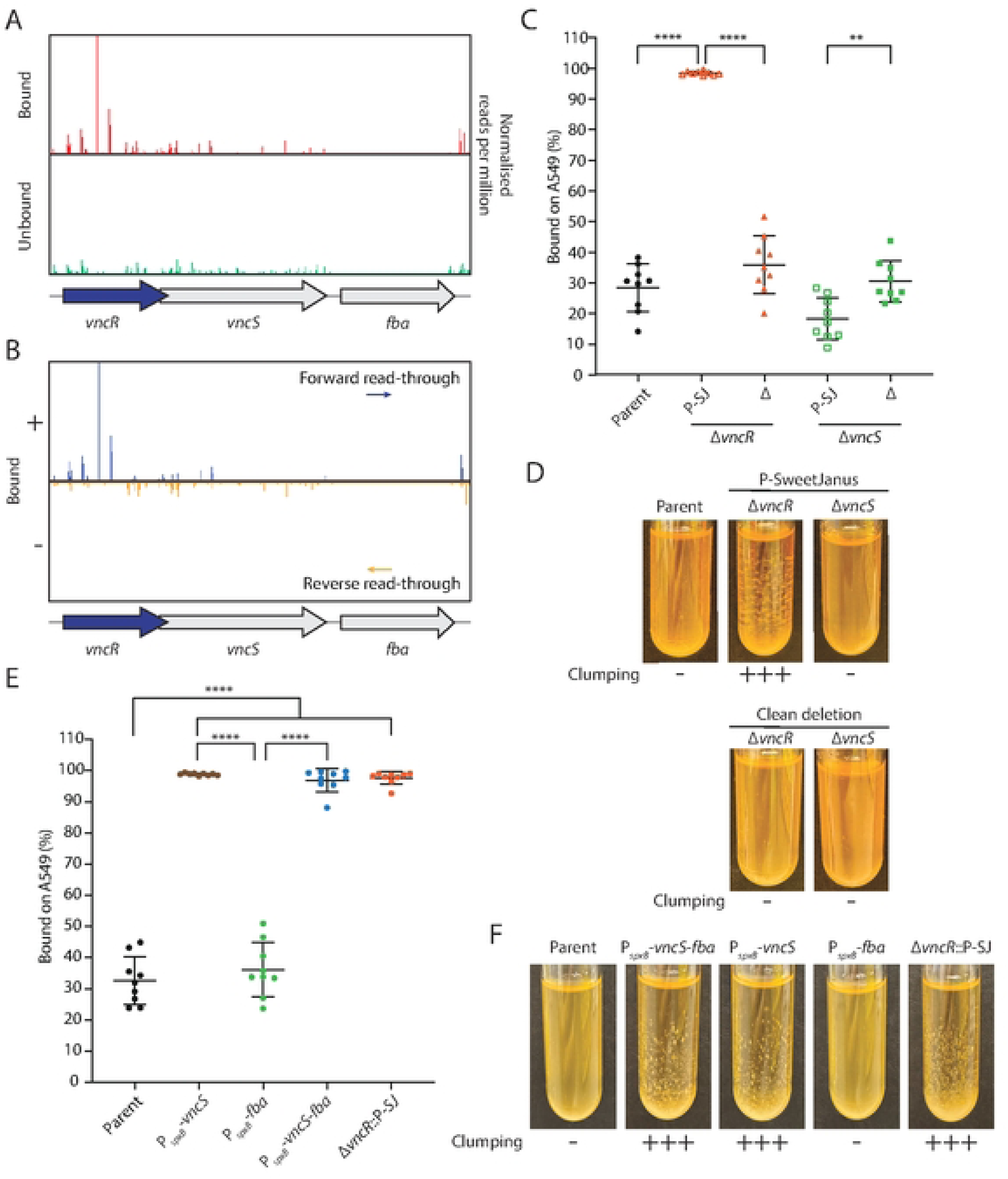
Overexpression of VncS, but not its cognate response regulator VncR, is required for elevated adhesion to A549 cells. (**A**) Genomic track showing normalized transposon insertion reads in the bound (top) and unbound (bottom) fractions at the *vncR–vncS–fba* locus. Insertions were enriched within *vncR* in the bound fraction, consistent with the Tn insertions being polar to the downstream *vncS*. (**B**) Strand-resolved insertion plot at the *vncR–vncS–fba* locus. Forward-strand insertions (blue, +) predominated in the bound fraction, suggesting transcription readthrough driving *vncS* overexpression. (**C**) Quantification of A549 cells adhesion for the indicated strains relative to their progenitor strain (HMS0002) after 3 hours of co-culturing. P-SJ insertion in *vncR* (NUS4970), but not *vncS* (NUS4973), significantly increased adhesion. Each data point represents an independent replicate. Bars show mean ± SD. **, *P* < 0.01; ****, *P* < 0.0001 by one-way ANOVA with Tukey’s post-hoc test. (**D**) Overnight cultures of P-SJ-disrupted and clean deletion strains. Clumping was observed only in Δ*vncR*::P-SJ, confirming that downstream *vncS* overexpression, not loss of *vncR*, caused the phenotype. (**E**) Quantification of A549 cells adhesion (% bound) for the indicated relative to their progenitor strain. Each data point represents an independent replicate. Bars show mean ± SD. ****, *P* < 0.0001 by one-way ANOVA with Tukey’s post-hoc test. (**F**) Overnight cultures of the indicated strains. The experiment was conducted three times with similar results.

Fba has also been reported to moonlight as a surface-exposed adhesin [33]. To directly examine the role of Fba in the increase in binding, we placed *vncS* and *fba* individually (P*_spxB_*-*vncS* and P*_spxB_*-*fba*) or in tandem (P*_spxB_*-*vncS-fba*) under the control of P*_spxB_*, a strong constitutive promoter of the *spxB* gene in *S. pneumoniae* [34,35]. To verify VncS overexpression, we inserted a FLAG tag at its C-terminus and demonstrated its functionality by showing that the FLAG-tagged strains retained the ability to form clumps in overnight cultures (**Fig. S4**). Immunoblotting confirmed that VncS was significantly overproduced in P*_spxB_-vncS* and P*_spxB_-vncS-fba* strains (**Fig. S4**). Under these conditions, P*_spxB_-vncS* and P*_spxB_-vncS-fba* displayed a ∼3-fold increase in A549 adhesion, comparable to the Δ*vncR*::P-SJ strain. Again, this result is consistent with the pronounced clumping in overnight cultures (**Fig. 3**). In contrast, the P*_spxB_-fba* strain neither increased adhesion nor clumping (**Fig. 3**). It is possible that, even under the merodiploid condition, Fba was not overexpressed in the P*_spxB_-fba* strain. To test this, we fused Fba with a C-terminal FLAG tag and found that Fba-FLAG levels did not increase significantly unless *fba* was co-transcribed with *vncS* (**Fig. S5**). This result indicates that *vncS* may be required for the stable accumulation of *fba* mRNA. Thus, we determined whether the kinase and phosphatase activities of VncS are required for the adhesion and clumping phenotypes in the P*_spxB_-vncS-fba* background. We introduced point mutations at the conserved catalytic residues, H241 (H241A, kinase-inactivated) and T245 (T245A, T245R, T245Y, phosphatase-impaired). Clumping was completely abolished in the P*_spxB_*-*vncS*^H241A^-*fba* and P*_spxB_*-*vncS*^T245Y^-*fba* strains and was markedly reduced in the P*_spxB_*-*vncS*^T245A^-*fba* and P*_spxB_*-*vncS*^T245R^-*fba* mutants relative to their parent strain (**Fig. S5**). Importantly, none of these mutations altered *fba* expression (**Fig. S5**), indicating that *fba* accumulation is unrelated to the clumping phenotype. We conclude that VncS activity is required for downstream signaling that promotes bacterial adhesion and that this signaling is likely independent of the cognate response regulator VncR and the presumed adhesin Fba.

### VncS crosstalks with RR06 to activate *cbpA* expression

Crosstalk between TCS pairs integrates multiple environmental signals through shared effector genes [36–38]. Having established that overexpression of VncS, but not disruption of VncR, led to elevated *cbpA* expression and increased adhesion, we hypothesized that VncS may crosstalk with a non-cognate response regulator. To test this hypothesis, we compared the transcriptomes of P*_spxB_-vncS-fba* and its progenitor strains (**Fig. 4**). This comparison revealed many differentially expressed genes in the P*_spxB_-vncS-fba* mutant, including downregulation of sugar transport genes (e.g., *spd_1675*) and carbohydrate metabolism enzymes such as an α-galactosidase (*spd_1678*) and a putative α-1,6-mannosidase (*spd_1972*), consistent with metabolic adaptation to *fba* overexpression. Furthermore, the *spd_2018-cbpA* locus was significantly upregulated (**Fig. 4**). qRT-PCR confirmed that *cbpA* transcript levels were elevated approximately 6-fold in P*_spxB_*-*vncS* relative to the progenitor strain (**Fig. 4**). To identify the specific transcriptional responses that cause the phenotype, we performed RNA sequencing of P*_spxB_-vncS-fba* and P*_spxB_-vncS*^H241A^-*fba* mutants (**Fig. 4**). Under these conditions, only the *spd_2018-cbpA* operon was differentially regulated (**Fig. 4**). The absence of other changes in this experiment confirms that the differential expression of metabolic genes was due to *fba* overexpression rather than VncS activity. Next, we deleted *cbpA*, *spd_2018*, or both in the P*_spxB_*-*vncS* background (**Fig. 4**). Deletion of *cbpA*, alone or in combination with *spd_2018*, completely abolished clumping in overnight cultures. Deletion of *spd_2018* did not affect clumping, indicating that *spd_2018* is not involved in the clumping phenotype. Since TCS06 (RR06/HK06) is known to regulate *cbpA* expression [20–22], we wondered whether VncS cross-phosphorylates RR06 and activates *cbpA*. To test this, we deleted *rr06*, *hk06*, or the entire TCS06 locus in the P*_spxB_*-*vncS* background (**Fig. 4C**). Deletion of *rr06*, with or without Δ*hk06*, abolished clumping. In contrast, deletion of *hk06* had a relatively minor effect, consistent with Δ*hk06* causing a slight elevation in *cbpA* expression (**Fig. 2**). *cbpA* transcript levels were also in line with the clumping results. Deletion of *rr06* in the P*_spxB_*-*vncS* background significantly reduced *cbpA* expression, whereas the kinase-defective VncS(H241A) and the phosphatase-defective VncS(T245Y) variants reduced *cbpA* expression to near wild-type levels (**Fig. 4**). We also assessed VncS variant expression by fusing a FLAG tag to the C-terminus. Again, none of these mutations altered VncS expression (**Fig. S6**). These results suggest that VncS activity in the P*_spxB_-vncS* cassette is required to phosphorylate RR06, and that RR06 phosphorylation is required for downstream activation of *cbpA*. Finally, to confirm that the VncS-RR06-*cbpA* axis translates into host cell adhesion, we measured A549 binding in strains carrying VncS kinase mutations or RR06 deletions in the P*_spxB_*-*vncS* background (**Fig. 4**). As expected, P*_spxB_*-*vncS*^H241A^ and Δ*rr06* // P*spxB*-*vncS* strains showed markedly reduced binding compared with the P*_spxB_*-*vncS* mutant, whereas deletion of *cbpA* in the P*_spxB_*-*vncS* background reduced A549 adhesion to near wild-type levels (**Fig. 4**). Thus, VncS is likely capable of cross-phosphorylating the non-cognate response regulator RR06 and activating *cbpA* expression. This activation may enhance pneumococcal adhesion to host respiratory cells.

**Figure 4.**
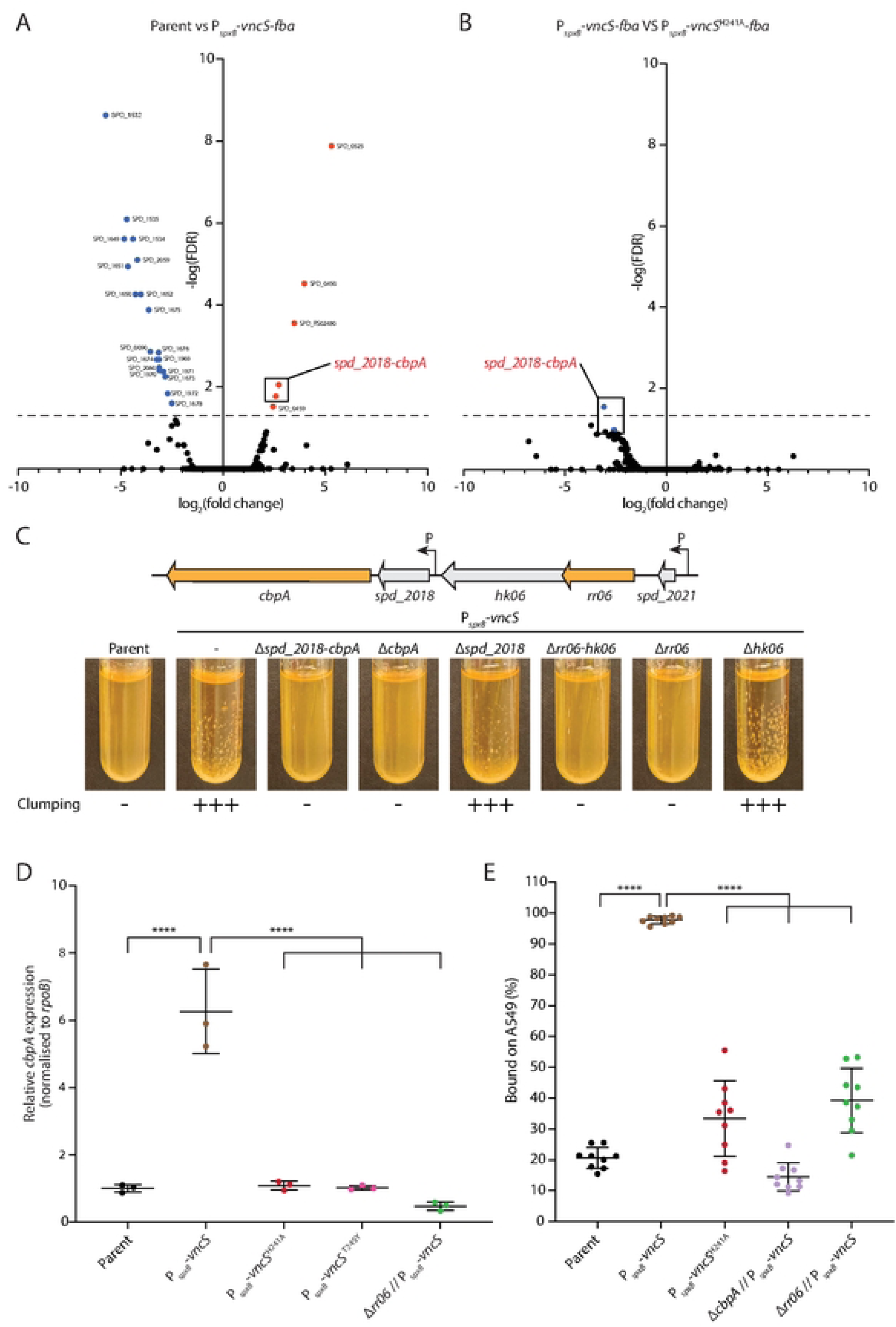
VncS activates *cbpA* expression through the non-cognate response regulator RR06. Volcano plots of the RNA sequencing data comparing gene expression (**A**) between P*_spxB_*-*vncS*-*fba* (NUS5242) and its progenitor strain (HMS0002), and (**B**) between P*_spxB_*-*vncS*-*fba* and P*_spxB_*-*vncS*^H241A^-*fba* (NUS5569). Genes significantly upregulated are highlighted in red, and those significantly downregulated are in blue. (**C**) Schematic of the *cbpA-spd_2018-hk06-rr06-spd_2021* locus (top) and overnight cultures (bottom) of the indicated deletion strains in the P*_spxB_*-*vncS* background. The experiment was conducted three times with similar results. (**D**) Relative *cbpA* transcript levels measured by qRT-PCR (normalized to *rpoB*) in strains P*_spxB_*-*vncS* (NUS5170), P*_spxB_*-*vncS*^H241A^ (NUS6896), P*_spxB_*-*vncS*^T245Y^ (NUS6899), and Δ*rr06* // P*_spxB_*-*vncS* (NUS6927). Each data point represents an independent replicate. Bars show mean ± SD. ****, *P* < 0.0001 by one-way ANOVA with Tukey’s post-hoc test. (**E**) Quantification of A549 cells adhesion (% bound) for strains P*_spxB_*-*vncS*, P*_spxB_*-*vncS*^H241A^, Δ*cbpA* // P*_spxB_*-*vncS* (NUS6926), and Δ*rr06* // P*_spxB_*-*vncS*. Loss of VncS kinase activity or deletion of *rr06* or *cbpA* reduces adhesion to near-progenitor levels. Each data point represents an independent replicate; Bars show mean ± SD. ****, *P* < 0.0001 by one-way ANOVA with Tukey’s post-hoc test.

### RR06 is cross-phosphorylated by VncS and dephosphorylated by HK06

We next asked whether phosphorylation of RR06 at its conserved receiver aspartate is required for this activation. First, we introduced phospho-ablative (D51A and D51N) and phospho-mimetic (D51E) substitutions in RR06 [39,40]. Then, we assessed whether these mutants clumped in the P*_spxB_*-*vncS* background (**Fig. S7**). Both phospho-ablative substitutions completely abolished clumping (**Fig. S7**). We measured the protein stability of these mutants and showed that RR06^D51A^ was produced at near-wild-type levels (**Fig. S7**), confirming that loss of clumping was likely due to an inability to receive the phosphoryl group at D51. By contrast, RR06^D51N^ was nearly undetectable by immunoblotting (**Fig. S7**), indicating severe protein instability that likely accounts for the absence of clumping. The phospho-mimetic RR06^D51E^ was stably expressed but failed to restore clumping (**Fig. S7**). These results are puzzling because both the phospho-mimetic and phospho-ablative mutants produced a similar phenotype, consistent with a previous report [21]. Nevertheless, these experiments confirm that phosphorylation of the D51 residue is required for clumping.

To define the relative contributions of VncS and HK06 to RR06 phosphorylation *in vivo*, we generated the Δ*vncS* Δ*hk06* double deletion and compared it with single deletions (**Fig. 5**). While Δ*hk06* produced weak but detectable clumping, the Δ*vncS* Δ*hk06* double deletion completely abolished clumping (**Fig. 5**). The Δ*hk06* Δ*cbpA* double mutants also did not clump, confirming that *cbpA* is required. qRT-PCR showed that *cbpA* levels in Δ*hk06* strains were mildly elevated relative to the progenitor strain, whereas in the Δ*vncS* Δ*hk06* mutant, *cbpA* levels were significantly reduced (**Fig. 5**). Thus, VncS likely provides the residual kinase activity that phosphorylates RR06 when HK06 is absent. Δ*vncS* alone had no significant effect on *cbpA* levels relative to its parent strain (**Fig. 5**), consistent with VncS operating below the detection threshold under laboratory conditions.

**Figure 5.**
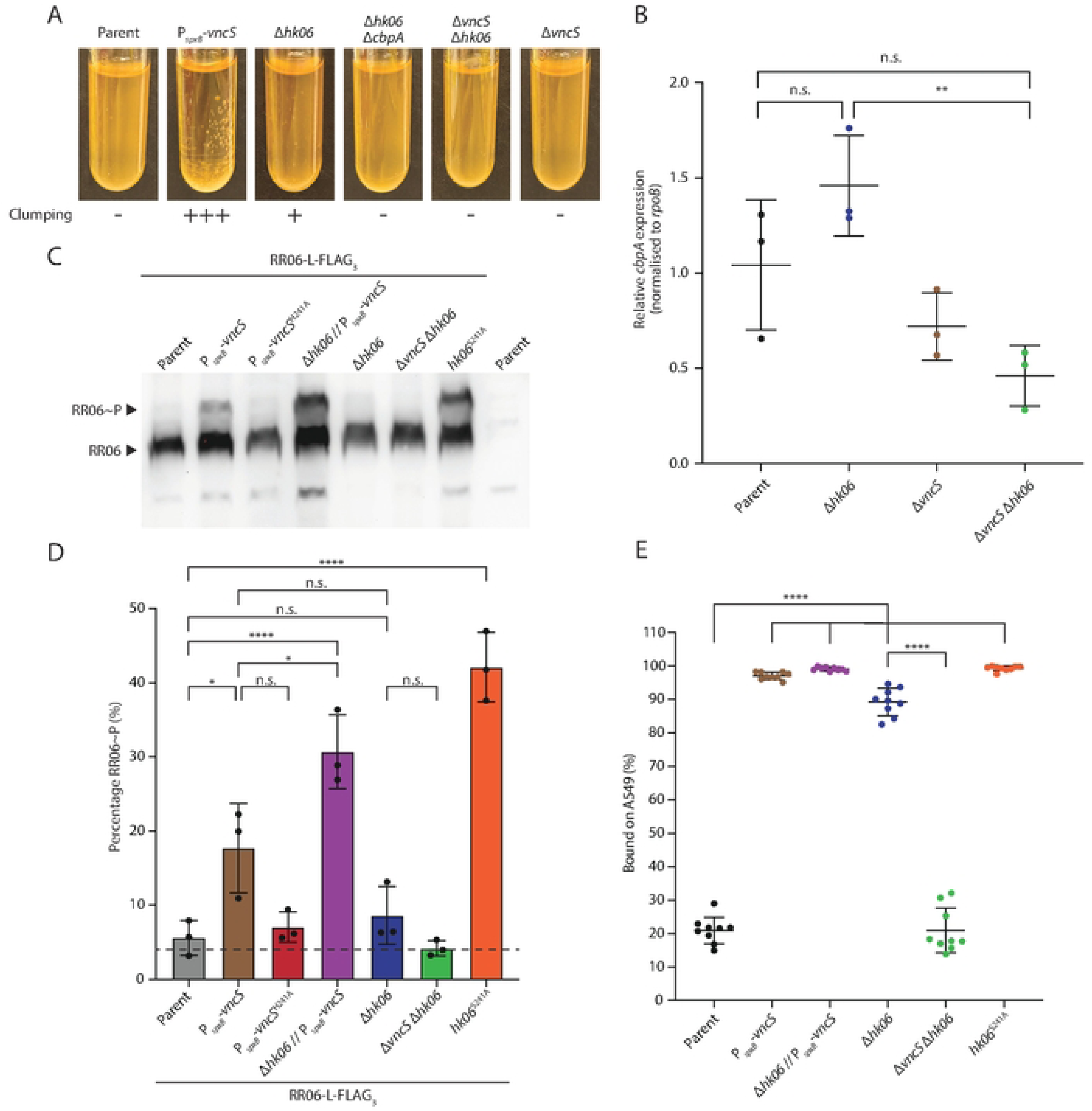
VncS phosphorylates RR06 *in vivo* and is controlled by the HK06 phosphatase activity. (**A**) Overnight culture of the progenitor strain (HMS0002), P*_spxB_*-*vncS*, Δ*hk06*, Δ*hk06* Δ*cbpA*, Δ*vncS* Δ*hk06* (NUS7189), and Δ*vncS* (NUS5008) grown in BHI broth for 14 to 18 hours at 37°C in 5% CO_2_. Clumping in Δ*hk06* is abolished by deletion of *cbpA* or *vncS*, demonstrating that residual *cbpA* activation in Δ*hk06* depends on VncS. (**B**) Relative *cbpA* transcript levels of the indicated strains measured by qRT-PCR (normalized to *rpoB*). Each data point represents an independent biological replicate. Bars show mean ± SD. **, *P* < 0.01; n.s., not significant by one-way ANOVA with Tukey’s post-hoc test. (**C**) Phosphorylation state of RR06 in various mutants. Strains harboring *rr06*-L-FLAG_3_ (NUS7390), *rr06*-L-FLAG_3_ // P*_spxB_*-*vncS* (NUS7573), *rr06*-L-FLAG_3_ // P*_spxB_*-*vncS*^H241A^ (NUS7574), *rr06*-L-FLAG_3_ Δ*hk06* // P*_spxB_*-*vncS* (NUS7670), *rr06*-L-FLAG_3_ Δ*hk06* (NUS7671), Δ*vncS rr06*-L-FLAG_3_ Δ*hk06* (NUS7672), *rr06*-L-FLAG_3_ *hk06*^S241A^ (NUS7391), and their progenitor strain (HMS0002) were grown to the early log phase in BHI broth. Cultures were normalized based on their optical densities. Cell lysates were resolved on an SDS-PAGE gel containing Phos-tag acrylamide, resolving phosphorylated (RR06∼P, upper band) from unphosphorylated (RR06, lower band) species. FLAG-tagged RR06 was detected by immunoblotting. Shown is a representative blot from three biological replicates. (**D**) Quantification of the percentage of RR06∼P relative to total RR06 from Phos-tag immunoblots in the indicated strains. Band intensities were determined by densitometry. The dashed line indicates the level in the Δ*vncS* Δ*hk06* strain. Each bar represents the mean ± SD of three biological replicates. Statistical analysis was performed using one-way ANOVA with Tukey’s post hoc test. *, *P < 0.05;* ****, *P* < 0.0001; n.s., not significant. (**E**) Quantification of A549 cells adhesion (% bound) in strains with different RR06 phosphorylation levels. Strains P*_spxB_*-*vncS*, Δ*hk06* P*_spxB_*-*vncS* (NUS7188), Δ*hk06*, Δ*vncS* Δ*hk06,* and *hk06*^S241A^ (NUS7220) were used to infect A549 cells. Each data point represents an independent replicate. Bars show mean ± SD. ****, *P* < 0.0001; n.s., not significant by one-way ANOVA with Tukey’s post-hoc test.

To dissect the catalytic activities of HK06, we introduced point mutations at its conserved histidine (H242A; primarily affecting kinase activity), threonine (T246A, T246R, T246Y; primarily affecting phosphatase activity), and serine (S241A and S241D; primarily affecting phosphatase activity) [22] (**Fig. S8**). Substitutions at H242 and T246 each produced modest clumping, indistinguishable from Δ*hk06*. Strikingly, the mutant harboring HK06^S241A^ exhibited a strong clumping phenotype that surpassed that of Δ*hk06*. qRT-PCR revealed that *cbpA* was upregulated by ∼13-fold relative to its progenitor strain, significantly higher than P*_spxB_-vncS* (∼4-fold) and Δ*hk06* (∼1.3-fold) (**Fig. S8**). On the other hand, the S241D substitution did not result in clumped cultures (**Fig. S8**), possibly because it does not affect phosphatase activity as strongly as S241A. Genetic epistasis analysis in the *hk06*^S241A^ background showed that deletion of *cbpA* or *rr06* abolished clumping, whereas deletion of *vncS* had no effect (**Fig. S8**). This demonstrates that HK06^S241A^ drives *cbpA* expression through RR06 and is independent of, or downstream of, VncS.

One reason RR06^D51A^ and RR06^D51E^ may have similar phenotypes is that glutamate may not substitute for the phosphorylated aspartate [22]. To directly measure RR06 phosphorylation, we employed Phos-tag acrylamide gel electrophoresis [41,42]. This approach resolves phosphorylated RR06∼P from unphosphorylated RR06 (**Fig. 5**). In the progenitor strain, RR06∼P was at a low level (∼5%), which increased to ∼18% in the P*_spxB_*-*vncS* mutant (**Fig. 5**). This increase was abolished by substituting VncS with the kinase-defective variant VncS^H241A^ (∼6%), demonstrating that VncS phosphorylates RR06 *in vivo* (**Fig. 5**). Consistent with the premise that crosstalk in TCS is controlled by the phosphatase activity of the cognate histidine kinase [36], deletion of *hk06* in the P*_spxB_*-*vncS* background (Δ*hk06* // P*_spxB_*-*vncS*) further elevated RR06∼P to ∼30% (**Fig. 5**). Deletion of *hk06* alone also modestly raised RR06∼P to ∼8%. In contrast, the Δ*vncS* Δ*hk06* double mutant reduced RR06∼P to ∼4%, reinforcing that VncS is responsible for RR06 phosphorylation in the absence of HK06. The phosphoaberrant *hk06*^S241A^ mutant has the highest RR06∼P level of all tested strains (∼42%), corroborating its high *cbpA* expression level and strong clumping phenotype (**Fig. S8**). Finally, A549 adhesion assays demonstrated that RR06 phosphorylation status directly correlates with host cell binding (**Fig. 5**). Strains with high RR06∼P: P*_spxB_*-*vncS*, Δ*hk06* // P*_spxB_*-*vncS*, and *hk06*^S241A^, all bound A549 cells at high levels (∼95–100%), whereas the Δ*vncS* Δ*hk06* mutant with near-basal RR06∼P adhered to A549 cells at levels comparable to their progenitor strain. We conclude that VncS can cross-phosphorylate RR06, increasing *cbpA* expression and adhesion to A549 cells. Meanwhile, HK06 mitigates crosstalk by acting as a phosphatase, thereby reducing RR06 phosphorylation and controlling its regulon.

## DISCUSSION

During infection, bacterial pathogens must continuously sense and adapt to the distinct microenvironments they encounter across diverse host niches. For *S. pneumoniae*, the transition from asymptomatic nasopharyngeal colonization to invasive disease requires adaptation from the mucosal surface to the bloodstream and, in the case of meningitis, to the central nervous system. This requires dynamic, coordinated regulation of surface virulence factors, particularly adhesins that mediate effective attachment to host epithelial cells [6,9]. In *S. pneumoniae*, virulence gene expression is regulated predominantly by two-component signal transduction systems (TCSs). The pneumococcal genome encodes 13 TCS pairs and one orphan response regulator [43,44], of which at least eight have been directly implicated in virulence [45]. For example, CiaRH senses sialic acids released by NanA to upregulate *nanA* expression, reinforcing colonization, while also regulating the serine protease HtrA to suppress the competence pathway [45–47]. Pan-genome-wide association studies have recently expanded the known regulons of multiple TCSs and identified novel response regulator-binding motifs [48]. Yet the full signaling cascade that controls adhesin expression in response to host signals has remained incompletely understood. Here, we used a barcoded Tn library screen to identify genetic determinants of adhesion to a human alveolar epithelial cell line. This work elucidated the regulatory mechanism controlling expression of the adhesin CbpA, a key mediator of nasopharyngeal attachment, complement evasion, and epithelial invasion [16–18]. Beyond the established role of TCS06 [20–22], we demonstrate that VncS can also control *cbpA* expression through cross-phosphorylation of the non-cognate response regulator RR06, and that HK06 functions primarily as a phosphatase that limits this crosstalk (**Fig. 6**). These findings also resolve a longstanding contradiction, where earlier work showed that both phospho-ablative and phospho-mimetic substitutions at RR06 abolished *cbpA* expression, leaving the mechanism of RR06 activation ambiguous [21,22]. Our Phos-tag data demonstrate directly that phosphorylation of RR06 is required for *cbpA* activation. VncS is the kinase responsible under the condition where VncS is overproduced or HK06 phosphatase activity is lost.

**Figure 6.**
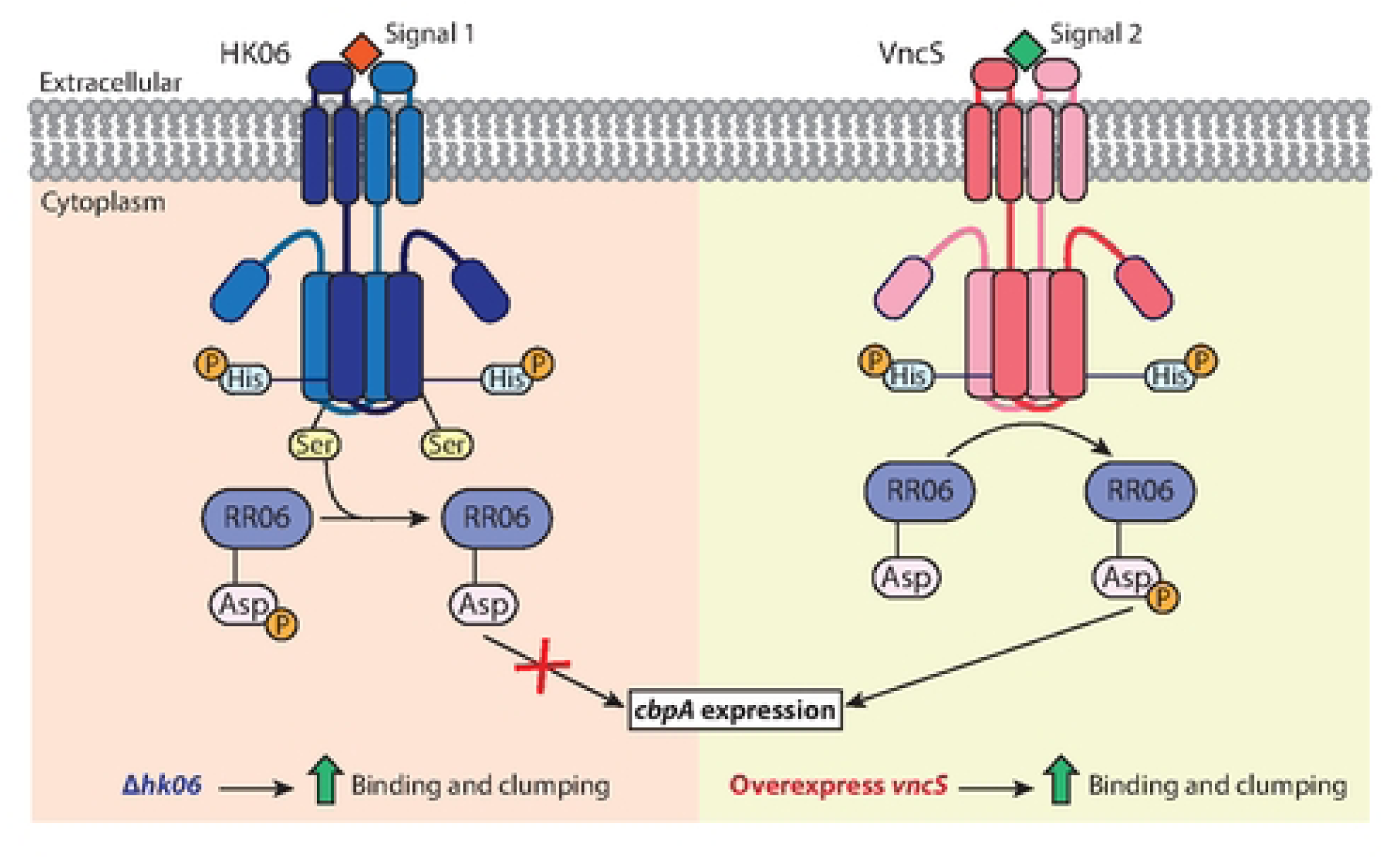
Model for VncS-RR06 crosstalk and HK06-mediated control of *cbpA* expression. Depicted is a model of the two-kinase regulatory circuit governing *cbpA* expression. Under basal conditions (left, pink background), HK06 senses an unknown signal via its extracellular sensor domain, autophosphorylates at the conserved His residue, and transfers the phosphoryl group to RR06 at the D51 residue to activate transcription of *cbpA*. Alternatively, HK06 functions as a phosphatase, mediated by its conserved S241 residue. This dephosphorylates RR06∼P, thereby limiting the duration and amplitude of *cbpA* induction. Loss of *hk06* abolishes this phosphatase activity, allowing residual RR06 phosphorylation by VncS to drive *cbpA* expression, which increases host-cell binding. Activation of VncS by another, yet-to-be-identified signal may activate RR06 and *cbpA* expression, a pattern mimicked by overexpressing *vncS*. This may counterbalance the control exerted by HK06 phosphatase activity, promoting bacterial adhesion in another host niche.

The physiological significance of inter-TCS crosstalk has long been debated, in part because it is typically observed only under conditions of gene deletion or overexpression [36]. Our unbiased transposon screen, however, revealed that not all overexpression events produce equivalent phenotypes. Among the 13 TCS histidine kinases in *S. pneumoniae*, only elevated VncS levels led to robust upregulation of *cbpA* and enhanced adhesion, indicating that this crosstalk is not generic but reflects a specific compatibility between VncS and RR06. This specificity raises the possibility that a physiological signal activating VncS would simultaneously engage the TCS06-*cbpA* circuit, enabling the pneumococcus to integrate distinct environmental inputs onto a shared regulon. A second layer of insulation against unwanted crosstalk is the stoichiometric imbalance between histidine kinases and their response regulators. In *E. coli*, the EnvZ-to-OmpR ratio has been estimated at approximately 1:30 [49], and overproduction of a non-cognate HK may overwhelm this buffer. Our data are consistent with this model because overproduction of VncS elevated RR06∼P and *cbpA* expression. This activation required not only the VncS kinase activity but was also amplified upon removal of the HK06 phosphatase, which keeps RR06 in its unphosphorylated, inactive state under standard laboratory conditions. These findings reinforce the general principle that the phosphatase activity of the cognate histidine kinase is the primary mechanism insulating TCS circuits from cross-activation [36,49]. It may also explain why the Δ*vncS* mutant is attenuated in vivo [50].

Several open questions warrant further investigation. First, overexpression of NanA active-site mutants did not abolish clumping, indicating that NanA generally increases surface adhesiveness rather than through its neuraminidase activity. Consistent with this, we could not detect the downstream glycosidases (BgaA, StrH, and EndoD) in the screen for factors involved in A549 cell attachment. Similarly, CbpA is known to be a pIgR-binding invasin [17]. However, A549 cells do not express pIgR. Still, CbpA overexpression robustly increases adhesion to A549 cells. Furthermore, the clumping phenotype suggests that bacterial cells adhere to one another in culture. These results indicate that CbpA and NanA function as general adhesins in this context. This is consistent with an earlier report that CbpA can facilitate nonspecific binding to host-cell surfaces independently of pIgR expression [51]. The structural basis for such adhesiveness remains unclear. Additionally, our screen did not detect all known adhesins, likely because their specific receptors were not expressed in A549 cells. Regarding HK06, our data reveal that the S241A variant, but not the T246A variant, elevates RR06∼P levels, suggesting that this residue is important for phosphatase activity. Mutations at the conserved Thr residue of the DHp domain may reduce both phosphatase and kinase activities, as evidenced by HK06 mutant phenotypes (**Fig. 5 and S8**). Finally, the physiological signal that activates VncS remains unknown. Lactoferrin activates VncS in vitro [50]. However, the required concentration is extraordinarily high (30 mM), far exceeding serum levels [50]. It is likely that VncS is induced by a serum-associated ligand distinct from lactoferrin, and once activated, VncRS may influence CbpA expression in addition to capsule production [52]. The latter response appears to be strain-specific [52]. Whether VncS is also activated in the nasopharyngeal cavity and the role of the potential VncS-RR06-CbpA axis in virulence will be addressed using appropriate infection models in the future.

## MATERIALS AND METHODS

### Bacterial strains and culture conditions

The strains used in this study are listed in Table **S1**. Unless otherwise specified, *S. pneumoniae* were grown in brain heart infusion (BHI) broth (Thermo Fisher Scientific, CM1135B), or on tryptic soy agar (TSA-II) supplemented with 5% (v/v) defibrillated sheep blood (blood agar; Biomed Diagnostics, 221261) at 37°C in 5% CO_2_. When indicated, antibiotics were added at a final concentration of 0.3 µg/mL for erythromycin (Erm), 300 µg/mL for kanamycin (Kan), 300 µg/mL for streptomycin (Str) and 150 µg/mL for spectinomycin (Spec).

### Strain construction

The primers used for generating PCR amplicons are listed in Table **S2**. Briefly, the PCR products were synthesized using Phusion DNA polymerase (New England Biolabs, M0530S) and purified using the QIAquick PCR purification kit (Qiagen, 28106). Briefly, *S. pneumoniae* strains were constructed by transforming cells with the isothermal assembly products or PCR amplicons after inducing natural competence with competence-stimulating peptide (CSP-1) for 9 minutes [53,54]. The cells were incubated at 37°C in 5% CO_2_ for 1.5 hours, and the transformants were plated on blood agar supplemented with either molten 0.7% (w/v) Difco nutrient agar (Biomed diagnostics, 213000) or molten 0.7% (w/v) Difco nutrient agar supplemented with 40% (w/v) sucrose agar (Fisher Scientific, S8600/60) and the appropriate antibiotics. The genetic constructs were validated by PCR using PowerPol (ABclonal, RK20719) and followed by Sanger sequencing. Allelic exchange was done using the Sweet Janus cassette (P-*sacB*-*kan*-*rpsL*^+^) or an erythromycin-resistant derivative of the Janus cassette (P-*erm*-*rpsL*^+^) [31,55].

### RBloxSpec library construction

Genomic DNA from a barcoded transposon library with 620,372 unique transposon insertions was extracted from library ML3 (an RBloxSpec library) [30] using the DNeasy blood and tissue kit (Qiagen, 69606). Strain HMS0002 [D39W *rpsL1* Δ*cpsE*] was grown in BHI broth to an OD_600_ between 0.1 and 0.2 and diluted to an OD_600_ of 0.1 in BHI broth before transforming with approximately 300 to 400 ng of the purified DNA. The cells were incubated according to the standard genetic transformation conditions described above. The transformants were plated on 19 blood agar plates supplemented with spectinomycin and incubated overnight at 37°C in 5% CO_2_, yielding approximately 1,040,000 transformants. The colonies on each plate were scraped, resuspended in 3 mL of BHI broth, and collected in a 50 mL Falcon tube. The final culture was further diluted 10X in BHI broth. Glycerol was supplemented to a final concentration of 15% (v/v), and the mixture was aliquoted and stored at -80°C.

### Epithelial cell adhesion assay

A549 cells (American Type Culture Collection, CCL-185) were cultured in Dulbecco’s Modified Eagle Medium (DMEM; Biowest, L0104-500) supplemented with 10% (v/v) fetal bovine serum (FBS; Biowest, S1810-500) and maintained at 37°C in 5% CO_2_. Unless otherwise specified, A549 cells were infected with *S. pneumoniae* strains as previously described [56]. Briefly, A549 cells were seeded overnight on 24-well plates (Corning Costar, CLS3526) at a density of 2.5 x 10^5^ cells per well. *S. pneumoniae* cultures were grown in BHI broth to an OD_600_ between 0.1 and 0.3 and normalized to an OD_600_ of 0.1 in DMEM. Bacterial infection was performed at a multiplicity of infection (MOI) of 10 (pneumococci to A549 cells). The co-culture was centrifuged at 300 x *g* for 5 minutes at 24°C and incubated at 37°C in 5% CO_2_ for 3 hours. After incubation, the supernatant containing unbound bacteria was collected, and the cells were washed three times with 1X phosphate-buffered saline (PBS; Vivantis, PB0344-1L). The cells with the bound bacteria were detached from the wells with 1X trypsin-ethylenediaminetetraacetic acid (EDTA) solution (Biowest, X0930-100) and quenched with an equal volume of DMEM supplemented with 10% FBS. The unbound and bound fractions were diluted in 1X PBS and plated on blood agar. Plates were incubated at 37°C in 5% CO_2_ overnight, and CFUs were enumerated. Infections using the transposon library were done in 6-well plates (Corning Costar, CLS3516) with A549 cells seeded at a density of 1.2 x 10^6^ cells per well and at an MOI of 10 (pneumococci to A549 cells).

### Barcode sequencing and quantification

After the infection experiments, genomic DNA of the ‘Input’, ‘Bound’, and ‘Unbound’ fractions of the barcoded isogenic capsule-switch mutants or the transposon libraries were extracted using DNeasy blood and tissue kit (Qiagen, 69606). Barcodes were amplified by the KAPA HiFi HotStart Library Amplification Kit (Roche, KK2611) using the primers listed in **Table S2**. The amplicons were gel-purified and sequenced using an Illumina platform (Azenta Life Sciences). The barcoded isogenic capsule-switch library was demultiplexed, and the raw barcode reads were enumerated using the CLC Genomics Workbench (Qiagen) as described in a previous study [28]. The reads of the ‘Input’, ‘Bound’, and ‘Unbound’ fractions from each serotype were normalized to the enumerated CFUs. The percentage of bound pneumococci in each experiment was calculated as the average of five separate wells from two batches of A549 cell cultures. The RBloxSpec library infection results were analyzed using published Perl scripts [30,57]. The barcode reads of each gene in the ‘Bound’ and ‘Unbound’ fractions were normalized to reads per million.

### Immunoblotting and Phos-tag analysis

Detection of FLAG-tagged proteins was done as described previously [53]. Briefly, strains expressing *vncS*-FLAG, *fba*-FLAG, and *rr06*-L-FLAG_3_ were grown in BHI broth till log phase and normalized to an OD_600_ of 0.3. Cells were collected by centrifugation at 8,000 x *g* for 1 minute and resuspended in 100 µL of 1X PBS. The cell suspension was mixed with 2X Laemmli buffer containing 5% (v/v) *β*-mercaptoethanol (BME; Bio-Rad, 16010710) at a 1:1 ratio and incubated at 50°C for 5 minutes. The samples were separated on a 12% SDS-PAGE gel and transferred to a polyvinylidene fluoride (PVDF) membrane (Bio-Rad, 1704272). The membrane was then blocked with 5% (w/v) skimmed milk in 1X PBST for 30 minutes, then incubated with rabbit anti-FLAG antibodies in blocking buffer (1:3,000 dilution; Sigma, F7425) overnight at 4°C. The membrane was washed twice with 1X PBST and incubated with goat anti-rabbit HRP antibodies (1:5,000 dilution; Invitrogen, A16110) for 1 hour at room temperature. The membrane was washed four times in 1X PBST before visualizing with enhanced chemiluminescence substrate (Thermo Fisher Scientific, 34580). The Phos-tag gel analysis was conducted as previously described [41,42]. Briefly, strains expressing *rr06*-L-FLAG_3_ were grown in BHI broth till log phase and normalized to an OD_600_ of 0.3. To collect the cells, 10 mL of culture was centrifuged at 8,000 x *g* for 5 minutes at 4°C, and the pellet was resuspended in 1 mL 20 mM Tris of pH 7.0 (Invitrogen, AM9850G) containing 8 µL protease inhibitor cocktail 3 (Merck, 539134-1MLCN) before transferring to a pre-chilled Lysing Matrix B tube (MP Biomedicals, 116911500). The bacteria were lysed using a Precellys Evolution homogenizer for three cycles at 6000 RPM for 40 seconds per cycle at room temperature, without pausing between cycles, and were placed on ice immediately. The lysate was centrifuged at 10,000 x *g* for 1 minute at 4°C, and the cell suspension was collected in an Eppendorf tube. Next, 36 µL of the sample was mixed with 4 µL of 5X Laemmli buffer containing 5% (v/v) *β*-mercaptoethanol, and separated on a 12% SDS-PAGE gel containing 75 µM Phos-tag acrylamide (Fujifilm, 304-93521). The samples were transferred onto a PVDF membrane, incubated with primary and secondary antibodies, and visualized as described above.

### RNA sequencing

Strains HMS0002 [D39W *rpsL1* Δ*cpsE*], NUS5242 [*rpsL1* Δ*cpsE* // ΔCEP::P*_spxB_*-*vncS*-*fba*], NUS5569 [*rpsL1* Δ*cpsE* // ΔCEP::P*_spxB_*-*vncS*^H241A^-*fba*-FLAG], NUS4966 [*rpsL1* Δ*cpsE Δhk06*] and NUS5169 [*rpsL1* Δ*cpsE Δhk06* // ΔCEP::P-*hk06*] were grown in BHI broth to an OD_600_ of 0.2 to 0.3 and normalized to OD_600_ of 0.2 in 1 mL. Next, 2 mL of RNAprotect Bacteria Reagent (Qiagen, 76104) was added to the culture, vortexed for 5 seconds, and then incubated for 5 minutes at room temperature. The culture was centrifuged at 5,000 x *g* for 10 minutes. The bacteria were subsequently lysed and RNA extracted using the RNeasy Mini kit (Qiagen, 74104) as described. Briefly, the pellet was resuspended in 200 µL of TE buffer supplemented with 600 µg/mL lysozyme (Gold Biotech, L-040-25), 15 µg/mL mutanolysin (Sigma-Aldrich, M9901), and 400 µg/mL proteinase K (Qiagen, RP107B). After incubating for 30 minutes at room temperature, 700 µL of RLT buffer supplemented with 1% (v/v) BME was added, and the mixture was vortexed vigorously. RNA was precipitated by adding 500 µL of absolute ethanol and transferred to a RNeasy spin column. The column was centrifuged at 10,000 x *g* for 30 seconds at room temperature and washed once with 350 µL of buffer RW1. DNA was removed by incubating for 15 minutes in 80 µL of DNase I incubation mix (Qiagen, 79254), followed by a final wash with 350 µL of buffer RW1. RNA was washed twice with buffer RPE and eluted with 50 µL of RNase-free water. The purified RNA samples were quantified by NanoDrop, and RNA sequencing was performed by Azenta Life Sciences.

### Quantitative real-time PCR (qRT-PCR)

RNA from various strains was extracted as described above. To quantify the expression level of *cbpA*, cDNA was synthesized from the RNA samples using the High-Capacity cDNA Reverse Transcription kit (Applied Biosystems, 4374966). Briefly, the RNA sample was diluted to 50 ng/µL in 10 µL. It was added to 2 µL of 10X RT buffer, 0.8 µL of 100 mM dNTP mix, 2 µL of 10X random primers, 1 µL of MultiScribe reverse transcriptase, 1 µL of RNase inhibitor, and nuclease-free water to a final volume of 20 µL. The samples were incubated in a thermocycler at 25°C for 10 minutes, 37°C for 2 hours, and 85°C for 5 minutes. The obtained cDNA was diluted 20X in water, and qPCR was performed using FastStart Essential DNA Green Master (Roche, 06402712001). For each reaction, 2 µL of diluted cDNA was added to 5 µL of master mix, 0.5 µL of the corresponding primer pairs listed in Table S2 at 10 µM, and 2 µL of water. The reactions were run on a LightCycler96 (Roche) with the following conditions: 95°C for 10 minutes; 45 cycles of 95°C for 10 seconds, 52°C for 10 seconds, 72°C for 10 seconds; and a final melting step of 95°C for 10 seconds, 65°C for 60 seconds, 97°C for 1 second. Relative *cbpA* and *pspA* expression were determined by the ΔΔCt method [58] and normalized to the relative expression level of *rpoB*.

### Data and statistical analysis

Unless otherwise stated, all statistical analyses were performed using GraphPad Prism 10 software. Non-parametric Spearman’s correlation was used to assess correlations in scatterplots. One-way ANOVA followed by Tukey’s multiple comparisons test was used to calculate P-values for qRT-PCR and A549 infection data.

## ACKNOWLEDGEMENTS

We thank Prof. Gan Yunn-Hwen for providing the A549 cells. We would also like to thank all members of the L.T.S. laboratory for the useful discussions.

## Funding

This work was supported by grants from:

The National Research Medical Council, Singapore (OFIRG23jan-0057 to LTS)

The Ministry of Education, Singapore (MOE-T2EP30220-0012 and MOE-T2EP30224-0012 to LTS)

The National Research Foundation (NRF-F-CRP-2024-0002 to LTS)

## Author Contributions

Conceptualization: S.-Y. T., Y.-Y. C., and L-T. S.

Methodology: S.-Y. T., J. J. Z., Y.-Y. C., K. S. T., and L-T. S.

Investigation: S.-Y. T., J. J. Z., and Y.-Y. C.

Visualization: S.-Y. T. and L-T. S.

Supervision: L.-T. S.

Writing – Original draft: S.-Y. T. and L-T. S.

Writing – review & editing: S.-Y. T., J. J. Z., Y.-Y. C., K. S. T., and L-T. S.

## Data and Materials Availability

All data are available in the main text or the supplementary materials.

